# Cooperation between Noncanonical Ras Network Mutations in Cancer

**DOI:** 10.1101/004846

**Authors:** Edward C. Stites, Paul C. Trampont, Lisa B. Haney, Scott F. Walk, Kodi S. Ravichandran

## Abstract

Cancer develops after the acquisition of a collection of mutations that together create the ‘cancer phenotype’. How collections of mutations work together within a cell, and whether there is selection for certain combinations of mutations, are not well understood. Using a Ras signaling network mathematical model we tested potential synergistic combinations within the Ras network. Intriguingly, our modeling, including a “*computational random mutagenesis”* approach, and subsequent experiments revealed that mutations of the tumor suppressor gene *NF1* can amplify the effects of mutations in multiple other components of the Ras pathway, including weakly activating, noncanonical, Ras mutants. Since conventional wisdom holds that mutations within the same pathway do not co-occur, it was surprising that modeling and experiments both suggested a functional benefit for co-occurring Ras pathway mutations. Furthermore, we analyzed >3900 sequenced cancer specimens from the Cancer Cell Line Encyclopedia (CCLE) and The Cancer Genome Atlas (TCGA) and we uncovered an increased rate of co-occurrence between mutations the model predicted could display synergy. Overall, these data suggest that selective combinations of Ras pathway mutations could serve the role of cancer “driver”. More generally, this work presents a mechanism by which the context created by one mutation influences the evolutionary trajectories of cancer development, and this work suggests that mutations that result in “network instability” may promote cancer in a manner analogous to genomic instability.

## INTRODUCTION

Cancer genomic studies support the idea that a cell becomes cancerous through the progressive acquisition of mutations that together confer the cancer phenotype. These mutations that promote cancer are commonly referred to as “driver genes” (Stratton et al., 2009). It is not well understood how the presence of one mutation might influence the selection of subsequent mutations through an evolutionary process (Yates and Campbell, 2012). Interestingly, there is statistical *lack* of co-occurrence between ‘canonical’ mutations within the same pathway (Thomas et al., 2007; Yates and Campbell, 2012). The lack of co-occurrence is typically attributed to the assumption that there would be no selective benefit to accumulating multiple mutations within the same molecular pathway (Yeang et al., 2008). Such arguments implicitly assume that each mutation is sufficiently strong to confer a selective advantage alone (e.g. the canonical *KRAS* and *BRAF* mutations). However, there are a number of weakly activating *RAS* mutations (Schubbert et al., 2006) and weakly activating *BRAF* mutations (Wan et al., 2004) that have been observed in cancer, although less commonly than the canonical mutations.

The role of noncanonical mutants in cancer is quite important when one considers the growing number of cancers that are being genomically characterized for both research and clinical purposes. It is believed that many cancers share common phenotypes, such as the constitutive activation of the Ras pathway (Hanahan and Weinberg, 2000). Within some types of cancer, there is near universal presence of a mutation that confers this phenotype to the Ras pathway. For example, essentially all sequenced pancreatic adenocarcinomas have canonical *KRAS* mutations (Biankin et al., 2012; Jones et al., 2008), and essentially all hairy cell leukemias have the canonical *BRAF* V600E mutation (Tiacci et al., 2011). More commonly, a type of cancer can utilize one of several potential gene mutations. Melanomas, for example, frequently harbor either a canonical *BRAF* or a canonical *NRAS* mutation (Hodis et al., 2012). When a canonical driver mutation is not identified in a sequenced cancer, other candidate driver mutations are often proposed based upon the identification of a mutated gene within the same pathway as a common, canonical, driver mutation (Hodis et al., 2012; Jones et al., 2008). Whether or not these less common mutations, which are often less strongly activating than the canonical mutations, are sufficient to serve as a surrogate for a canonical driver mutation, or whether the ability to serve as a surrogate is conditional to some other contextual influence, is not fully understood.

We set out to investigate oncogenic signaling within Ras network proteins and the potential for cooperation between less commonly mutated genes. We used a mathematical model to investigate whether canonical and noncanonical Ras mutants are influenced by the partial loss of tumor suppressor gene product neurofibromin (NF1). We found computational evidence for greater than additive increases in Ras activation for noncanonical Ras mutants within in the neurofibromin deficient context. This prediction was also supported experimentally in cells with or without neurofibromin. Further, analyzing recently available large cancer genomic data sets, we found an increase in coincidence between *NF1* mutations and mutations in other players within the Ras network. This work suggests that *NF1* mutations promote cancer not only by the direct and immediate increase in RasGTP, but also by increasing the number of possible subsequent mutations within the Ras pathway that would further increase Ras signaling. This finding contrasts conventional wisdom that mutations within the same pathway do not co-occur, and rather suggests that multiple, weakly activating mutations may together serve the role of driver gene. Overall, this work demonstrates how a biochemical/mechanistic understanding of cell signaling networks can be used to uncover functional combinations of mutations that promote cancer.

## RESULTS

### Modeling predicts synergy between weakly activating Ras pathway mutants

We previously developed a mathematical model based upon the biochemical reactions for the major classes of proteins that regulate Ras GTPase signaling (Stites et al., 2007). We had used this model to study the constitutively elevated levels of Ras pathway signals that are produced by canonical oncogenic Ras mutants. Among the predictions of our earlier model was that oncogenic Ras mutants drive the activation of wild-type Ras within the same cell. The different roles of wild-type and oncogenic mutant Ras in cancer have since become an important topic in cancer biology (Grabocka et al., 2014; Jeng et al., 2012; Lim et al., 2008; Young et al., 2013). The model also predicted the counterintuitive activation pattern of a Ras mutant that retained GTPase activity but was GAP-insensitive, and the later discovery of such a mutant in Noonan syndrome confirmed the predicted behaviors (Schubbert et al., 2007).

When we originally developed our Ras model, we focused upon neurofibromin as the basally active Ras GAP that maintains low-levels of Ras signal in non-stimulated cells (Ahmadian et al., 1997). Of all Ras GAPs, neurofibromin has the best evidence for contributing to the basal regulation of steady-state RasGTP levels and the clearest association with cancer. *NF1*, the gene coding for neurofibromin, is one of the most commonly mutated genes in lung cancer (Ding et al., 2008), ovarian cancer (Kan et al., 2010; The Cancer Genome Atlas Netwrok, 2011), and glioblastoma (Brennan et al., 2013; Parsons et al., 2008; The Cancer Genome Atlas Network, 2008). Loss of a single copy of tumor suppressor gene *NF1* does not appear capable of promoting cancer alone, although *NF1* displays haploinsufficiency and a loss of one functional allele is sufficient to cause a small increase in RasGTP and the cellular proliferation rate (Shapira et al., 2007). The disease neurofibromatosis is caused by germline loss of one functional copy of *NF1*, which further highlights that a small, partial increase in Ras signal can have pathological consequences. Individuals with neurofibromatosis also have an increased risk of developing many different tumors (Williams et al., 2009).

Here, we used this model to investigate the effect of concurrent Ras and neurofibromin mutations. Pathological *NF1* mutations, such as those associated with cancer and neurofibromatosis, commonly result in loss of expression of neurofibromin (The Cancer Genome Atlas Network, 2008). Gene mutations within the Ras pathway can also result in a protein product with altered biochemistry. For example, the oncogenic Ras mutant RasG12V has an apparent complete loss of k_cat_ for GAP activity on Ras, an order of magnitude decrease in its intrinsic GTPase reaction, slight variations in nucleotide affinity, and slightly increased affinity for its effector Raf. Such changes can be applied to the model to make predictions that match well with experimental data (Stites et al., 2007). Thus, mathematical models can find behaviors that naturally emerge from the changes in rate constants and concentrations that follow from gene mutation, and in that manner predict how the system responds to particular mutations (Stites and Ravichandran, 2012b).

We initially considered the canonical oncogenic Ras mutants RasG12D and RasG12V, and the noncanonical, weakly activating mutation RasF28L. When neurofibromin is fully present and not mutated, the RasF28L mutation was predicted to result only in approximately half the RasGTP signal as RasG12D or RasG12V mutation, which is consistent with experimental data for these mutations (Stites et al., 2007). However, when we modeled RasF28L in the neurofibromin deficient context, we observed that RasF28L generated a similar, high, level of Ras activation similar to the strong RasG12D and RasG12V mutations (Figure 1A, upper). This is due, in part, to a less-than-additive increase in Ras signals when RasG12D and RasG12V are combined (Figure 1A, lower). This less-than-additive increase may be consistent with the general lack of co-occurrence between commonly observed ‘strong’ mutations in that it suggests there is a smaller benefit to acquiring both mutations than is provided by either alone. Interestingly, RasF28L exhibited a greater than additive increase in Ras signals when combined with a loss of NF1 activity. We hypothesized that there may be a large number of such ‘inherently weaker’ Ras mutations beyond RasF28L that would result in high levels of Ras pathway activation in the neurofibromin deficient (NF1-deficient) conditions, but not in neurofibromin wild-type (NF1-WT) conditions.

**Figure 1.**
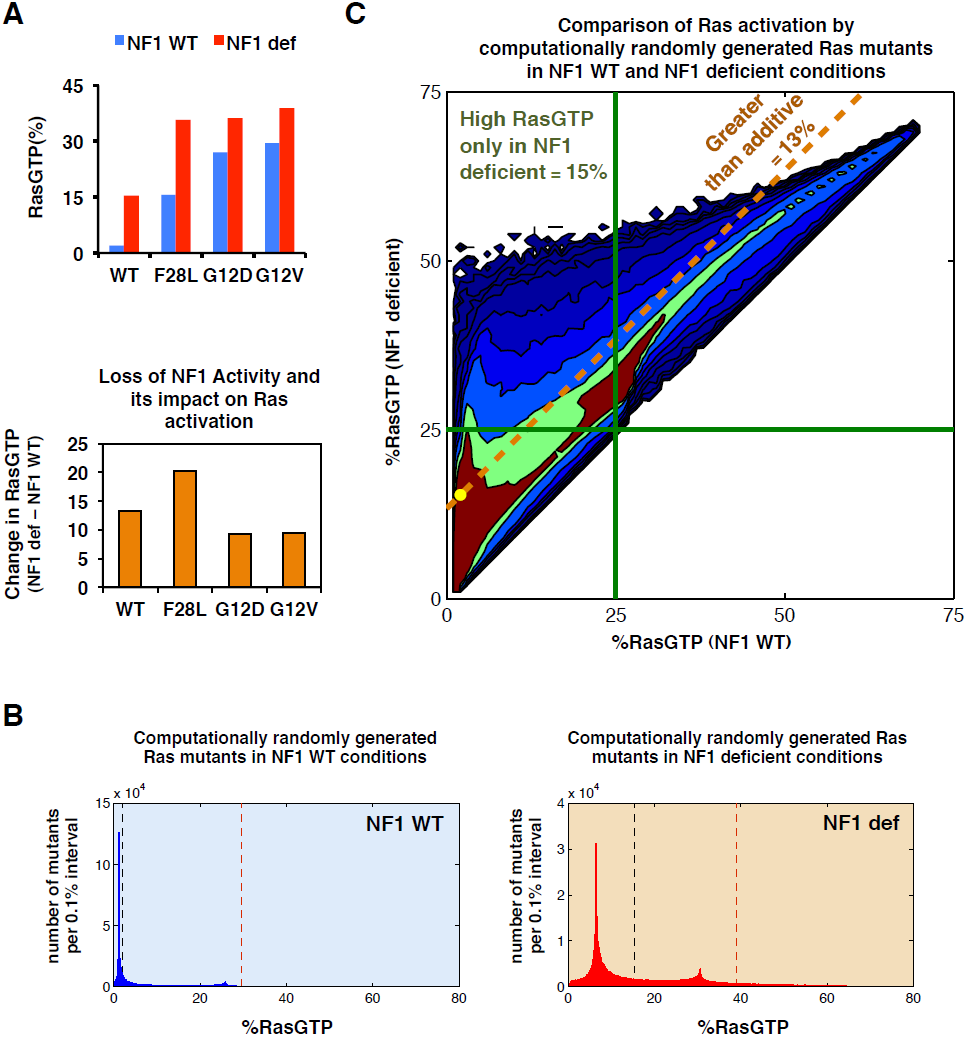
Modeling predicts that weakly activating Ras mutations can appear strong within the NF1-deficient context. A. (upper) Simulations of the Ras network model to evaluate the co-occurrence of an *NF1* loss-of-expression mutation with a canonical oncogenic mutation (RasG12D and RasG12V) or a noncanonical, non-oncogenic mutation (RasF28L). Conditions with no *RAS* mutation and no *NF1* mutation are also included. (lower) The net change in RasGTP levels predicted for going from NF1 wild-type to NF1-deficient conditions for the indicated Ras mutants compared to the same change in a Ras WT background.
B. Histogram from our ‘computational mutagenesis’ displaying the number of Ras mutants with varying levels of RasGTP signal in the context of NF1-WT (blue) and NF1-deficient (red) conditions. Dashed black line shows level of RasGTP for a network with all RasWT (no mutation present); dashed brown line shows level of RasGTP for a network with a RasG12V mutation. Histogram is binned into 0.1% intervals.
C. One million random mutants were simulated in the NF1-WT and NF1-deficient states, and the resulting levels of RasGTP are plotted for both conditions. The yellow dot marks RasGTP levels for WT and NF1-deficient networks in a network with only RasWT. Any random Ras mutant falling above the dashed gold line shows a greater net change in percent RasGTP in the NF1-deficient network compared to the WT network. Green lines indicate 25% total RasGTP.

### ‘Computational mutagenesis’ predicts that some weak Ras mutants could become strong in a neurofibromin deficient context

Very few Ras mutants have been characterized as extensively as the RasG12D and RasG12V mutants. We therefore performed a “computational random mutagenesis” to more comprehensively explore the possible extent to which neurofibromin deficiency might affect potential Ras mutations (see Methods and Supplemental Information). By varying the biochemical rate constants and enzymatic parameters that characterize a mutant Ras, and by simulating the Ras network under NF1-WT and deficient conditions, we investigated how variants of Ras that arise from this computational mutagenesis might result in altered Ras signaling. The key advantages of this computational random mutagenesis approach is the ability to investigate the behavior of mutants that might not confer a selective advantage (on their own), thereby enabling them to be observed in a human cancer sample, and the ability to rapidly investigate a wide range of mutants with widely varying properties. We simulated one million such random mutants and compared the RasGTP levels in the context of both the NF1-WT and NF1-deficient networks.

We analyzed the relative frequencies with which these randomly generated Ras mutants achieved different RasGTP levels (Figure 1B), and observed that, in the NF1-WT context, most spontaneous mutations did not cause a large increase in Ras signaling. A long, shallow tail of more strongly activating mutants was also noticed. This suggests that there are a limited number of potential Ras mutations that result in high levels of ‘constitutive’ Ras activation and are strong enough to promote cancer by themselves. In contrast, in the NF1-deficient context, the distribution was shifted toward higher levels of RasGTP, and the relative frequencies of mutants generating higher levels of RasGTP also increased. This could be interpreted to mean that the number of potential Ras point mutations capable of promoting cancer development would be greater in the NF1-deficient context.

We then examined how the level of RasGTP for each mutant differed between NF1-WT and NF1-deficient contexts, and we considered the potential synergy between neurofibromin and Ras mutants (Figure 1C). Approximately 13% of all random mutants displayed greater-than-additive increase in RasGTP under neurofibromin deficiency (i.e. the area above the orange diagonal line). We then focused on mutants that resulted in 25% of total Ras bound to GTP, since ∼25% of cellular RasGTP has previously been shown to approximate where transformation potential abruptly begins (Donovan et al., 2002). We found that 15% of all random mutants exceeded this 25% level of RasGTP, but only in the NF1-deficient context. The model thus predicts that the number of Ras mutations that could result in a high level of RasGTP will be increased in NF1-deficient conditions. Strikingly, these simulations also suggest that the strength of a mutant can be context dependent, with a Ras mutation that appears “weak” in a wild-type background being “strong” in an NF1-deficient background. More generally, our simulations suggest that the NF1-deficient conditions may make the context ‘permissive’ for the development of cancer by increasing the number of potential mutations that will sufficiently elevate RasGTP, and also increasing the probability that a randomly acquired mutation might become sufficiently strong to promote cancer.

### Ras F28L mutant is more strongly activating in the NF1-deficient cellular context

We next experimentally tested our computational prediction that some “weak” Ras mutants will more strongly activate the Ras pathway within the NF1-deficient context. We obtained mouse embryo fibroblasts (MEFs) that had been derived from *Nf1* deficient mice as well as the wild-type, and heterozygous null mice (*Nf1*-/- and *Nf1*+/+, *Nf1*+/-, respectively) (Shapira et al., 2007). We confirmed that the *Nf1*+/- and *Nf1*-/- MEFs had decreased protein expression (Figure 2A) and decreased mRNA expression for NF1 (Figure S1). The progressive decreases in neurofibromin expression level phenotypically resulted in increased rates of cellular proliferation (Figure S2). Readouts of RasGTP signaling, such as phosphorylated ERK (pERK) by Western blot (Figure 2A) or by flow cytometry (Figure 2B), reflected the different levels of neurofibromin deficiency. The modest changes to the expression of other GAPs were not sufficient to make up for the loss of neurofibromin GAP activity, as evidenced by the increased levels of RasGTP and pERK in *Nf1+/-* and *Nf1-/-* MEFs (Figures 2A and S1). These observations suggest that neurofibromin plays a critical and dominant role in these MEF cells, and that these MEFs could be used for our further analysis. Additionally, these observations experimentally reiterated that graded changes in Ras pathway activation have phenotypic consequences.

**Figure 2.**
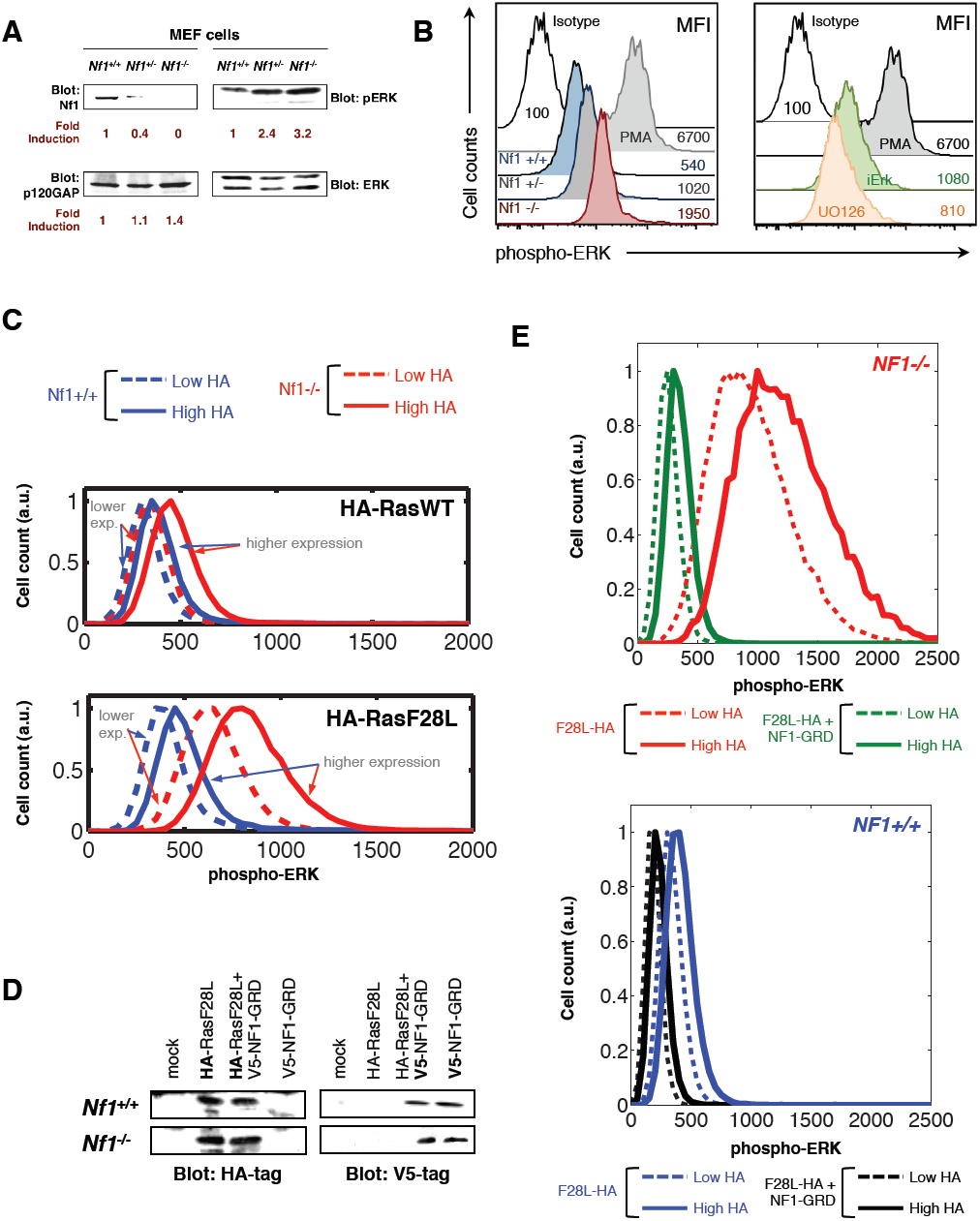
Weak Ras mutants in mammalian cells behave as strong activators of Ras pathway signaling under the *Nf1* deficient conditions. A. Immunoblots of *Nf1*+/+, *Nf1*+/-, and *Nf1*-/- mouse embryo fibroblasts (MEFs) for expression of neurofibromin, p120 Ras GAP, and phosphorylated ERK. Loss of neurofibromin expression leads to increased Ras pathway activation, as measured by phosphorylated ERK.
B. (left) Histograms present p-ERK profiles within control (*Nf1*+/+), *Nf1*-heterozygote (*Nf1*+/-) and *Nf1*-deficient (*Nf1*-/-) MEF cells. Mean fluorescence intensity for each condition is indicated on the right of histogram plots. Stimulation of cells with PMA served as a positive control for maximal pERK activation. Data presented are representative of at least 6 similar experiments. (right) The inhibition of the pERK signal inhibitory ERK peptide and the U0126 drug served to confirm the veracity of the observed pERK signals in this flow cytometry based detection assay.
C. *Nf1+/+* and *Nf1-/-* MEFs transfected with HA-tagged RasWT or HA-tagged RasF28L were divided into lower and higher HA-tag expressing populations by multi-color flow cytometry, where we gated on the signal for HA signal first and then plotted the pERK signal within these populations.
D. Immunoblots showing expression of HA-tagged RasF28L and V5-tagged NF1-GRD in *Nf1-/-* and *Nf1+/+* MEF cells following transfection, either alone or together.
E. Flow cytometry was used to gate on MEFs transfected with HA-tagged RasF28L or HA-tagged RasF28L and NF1-GRD into lower and higher HA-tag expressing populations, and the pERK signal within these populations is shown. *Nf1*(-/-) + F28L, red; *Nf1*(-/-) + F28L + NF1-GRD, green; *Nf1*(+/+) + F28L, blue; *Nf1*(+/+) + F28L + NF1-GRD, black. Higher HA, solid; lower HA, dashed.

We next examined how variation in the level of RasF28L expressed could affect ERK phosphorylation in *Nf1*+/+ and *Nf1*-/- conditions (since ERK activation is well established as one downstream readout of Ras pathway activation). We first transiently transfected HA-tagged RasF28L or HA-tagged RasWT into *Nf1+/+* and *Nf1-/-* MEFs. Flow-cytometry was used to obtain quantitative measurements of both the amount of transfected protein expression at a single cell level, and the amount of resulting Ras pathway activation within each cell. These measurements showed that RasF28L expression caused a larger increase in phosphorylated ERK signal in the *Nf1*-/- MEFs than it caused in the *Nf1*+/+ MEF (Figure 2C). Furthermore, gating the RasF28L cells based on relatively lower and relatively higher expression demonstrated that the change in ERK phosphorylation going from lower to higher RasF28L expression was concomitantly enhanced in the *Nf1*-/- MEFs than in the *Nf1*+/+ MEFs, consistent with the model predictions.

If the greater changes in phosphorylated ERK within *Nf1*-/- conditions were due to the loss of GAP activity by neurofibromin, then reintroduction of neurofibromin would be expected to reverse the increased change in RasGTP signal observed in *Nf1*-/- MEFs. We chose to express the GAP-related domain of neurofibromin (NF1-GRD), as this domain alone is sufficient to catalyze the hydrolysis of GTP on Ras. We co-transfected *Nf1*-/- and *Nf1*+/+ mouse embryo fibroblasts with HA-tagged RasF28L and with V5-tagged NF1-GRD (Figure 2D). We compared cells expressing lower and higher levels of HA-Ras F28L via the flow cytometry-based ERK activation assay. Co-expression of NF1-GRD reversed the large magnitude changes in pERK due to RasF28L expression in *Nf1*-/- MEFs (Figure 2E, upper). Furthermore, co-expression of NF1-GRD with RasF28L in wild type *Nf1*+/+ MEFs had a similar but smaller effect on changing the phosphorylated ERK signal as a function of RasF28L (Figure 2E, lower). The NF1-GRD experiments confirmed that the increased strength of F28L is due to differences in the Ras GAP activity of NF1. Overall, these experimental observations in mammalian cells support the important and non-obvious prediction from the mathematical model that neurofibromin deficiency can cause some normally “weak” Ras mutants to have a stronger effect on Ras pathway activation.

### Co-occurrence between NF1 mutations and noncanonical Ras mutations in cancer genomic data sets

We hypothesized that synergy between NF1 mutations and noncanonical Ras mutations is functionally important in some cancers, and that an increased co-occurrence of *NF1* and noncanonical *RAS* mutations should therefore be observable in cancer genome data sets. As noncanonical *RAS* mutations are much less commonly observed than canonical *RAS* mutations, an analysis of increased co-occurrence requires a large dataset. We first considered the Cancer Cell Line Encyclopedia (CCLE), which includes massively parallel sequencing data for greater than nine hundred cancer cell lines (Barretina et al., 2012). We investigated the frequency with which *NF1* mutations co-occurred with noncanonical *RAS* mutations (e.g. *KRAS*, *NRAS*, and *HRAS* mutations that are not at codon 12, 13, or 61) and with canonical *RAS* mutations (codon 12, 13, or 61) (Figure 3A, upper). We found that *NF1* mutations were much more common in cells that also harbored a noncanonical *RAS* mutation. There was a consistent increase for all three *RAS* isoforms, and the difference was statistically significant for *KRAS* (p<0.005 by Fisher’s Exact test). We compared the co-occurrence of mutations to *TP53* as a control, and we found no significant differences in the rate of co-occurrence (Figure 3A, lower). We note that although there was a similar increase of co-occurrence between noncanonical *NRAS* mutations with *NF1* mutations, that the CCLE dataset includes fewer noncanonical *NRAS* mutations (eleven) than noncanonical *KRAS* mutations (twenty-nine); the p-value for the difference (p=.06 by Fisher’s Exact test) was not quite significant due to the smaller number of cases. We highlight that even though we are examining the same number of total cancer genomes for the different combinations of gene mutations here and in the remainder of the manuscript, that the number of times a specific gene is found to be mutated can vary widely. Due to this variability, we present the percentage of the mutated gene of interest that co-occurs with another mutated gene (i.e. 10% of canonical KRAS mutations co-occur with an NF1 mutation) to facilitate comparisons between members of a class of genes, and use the p-value from the Fisher’s Exact test, which takes into account the exact number of mutations, genomes analyzed, and the various co-occurrences, to determine whether the pattern of co-occurrence is statistically significant.

**Figure 3.**
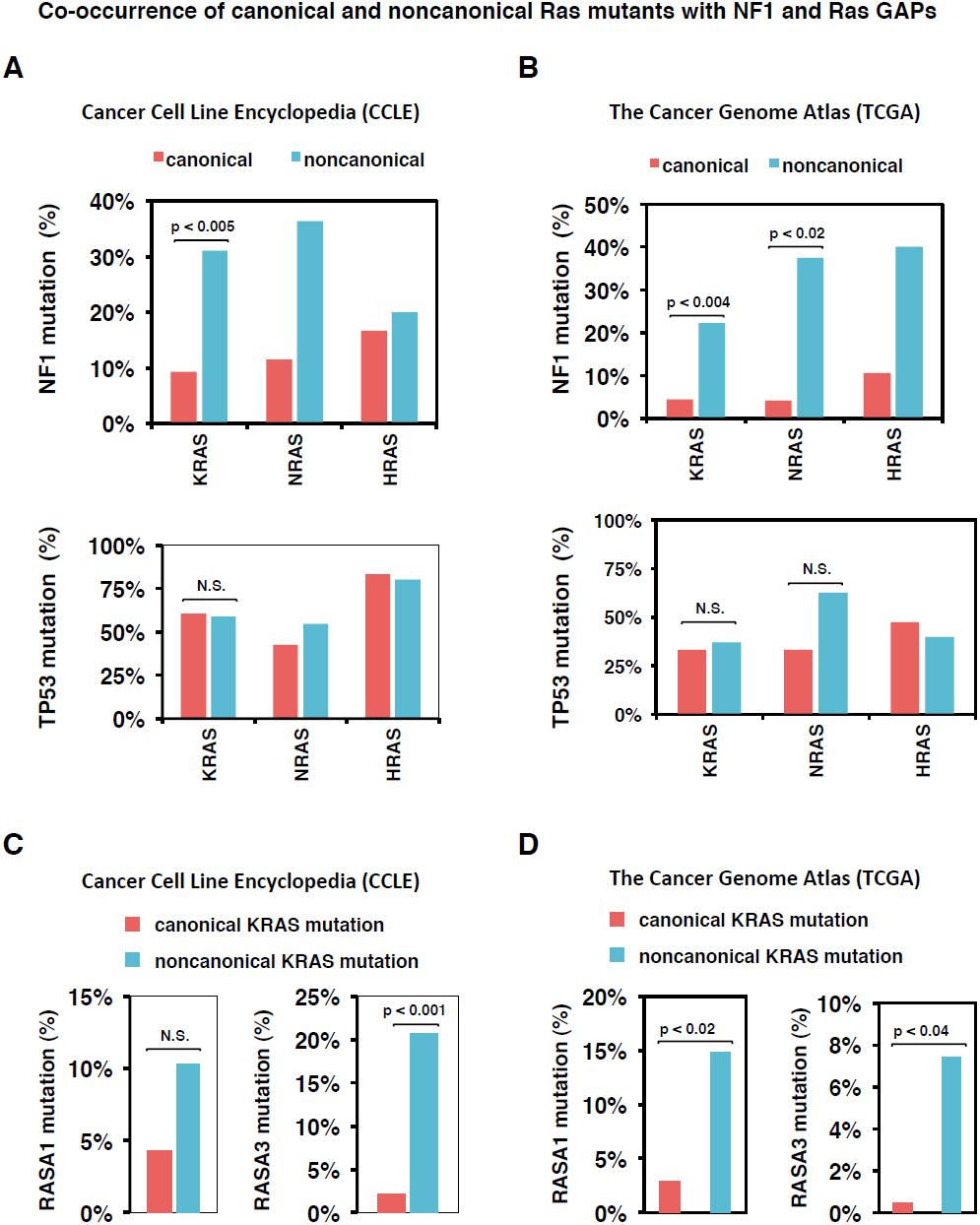
Co-occurrence of noncanonical Ras mutants with *NF1* and Ras GAP mutants. A. Percentage of canonical and noncanonical *KRAS*, *NRAS*, and *HRAS* mutants that co-occur with an *NF1* mutation (above) or with a *TP53* mutation (below) within the CCLE data on 947 cancer genomes (Barretina et al., 2012). The p values are indicated. N.S. refers to statistical differences that were not significant.
B. Percentage of canonical and noncanonical *KRAS*, *NRAS*, and *HRAS* mutants that co-occur with an *NF1* mutation (above) or with a *TP53* mutation (below) within the TCGA dataset of 3281 cancer genomes (Kandoth et al., 2013).
C. Percentage of canonical and noncanonical *KRAS* mutants that co-occur with a Ras GAP mutation within the CCLE dataset.
D. Percentage of canonical and noncanonical K-Ras mutants that co-occur with a *RASA1* or *RASA3* mutation within the TCGA data. The p-value for all panels is by Fisher’s exact test.

We next analyzed the recently published dataset from The Cancer Genome Atlas (TCGA), with whole exome and/or whole genome sequencing for more than three thousand different human cancer samples (Kandoth et al., 2013), for the proportion of canonical and noncanonical *KRAS*, *NRAS*, and *HRAS* mutations. Noncanonical *KRAS*, *NRAS*, and *HRAS* mutations were found to co-occur much more frequently with an *NF1* mutation (Figure 3B, upper). The increase was statistically significant for *KRAS* and *NRAS* (p<0.004 and p<0.02, respectively, by Fisher’s Exact test). Analysis of the rate of co-occurrence with *TP53* mutations as a control again found no significant differences in the rate of co-occurrence (Figure 3B, lower).

Thus, in two separately acquired, large, cancer data sets, there was evidence for an increased rate of co-occurrence between noncanonical *RAS* mutations and *NF1* mutations. Overall, the increased co-occurrence of *NF1* mutations with noncanonical *RAS* mutations is consistent with our prediction that the *NF1* mutation enables a larger number of less potent *RAS* mutations to have large, cancer promoting effects.

### Noncanonical Ras mutations are more likely to be found with Ras GAP mutations

Although we have focused on neurofibromin as the basally active Ras GAP, other Ras GAPs may also serve to maintain low levels of Ras signaling in non-stimulated conditions. For example, DAB2IP has been shown to serve the role of a basally active Ras GAP in prostate cancer cells (Min et al., 2010). As our model more generally describes the behavior of basally active Ras GAPs on regulating Ras pathway activation, we considered the possibility that other mutations to other Ras GAPs may similarly promote the effects of noncanonical Ras mutations.

We considered noncanonical *KRAS* mutations in both the CCLE data and TCGA data for co-occurrence with non-*NF1* Ras GAP mutations. Within the CCLE data set, we find increased rate of co-occurrence with Ras GAPs *RASA1* and *RASA3* for noncanonical *KRAS* mutations (Figure 3C). This difference was statistically significant for *RASA3* (p<0.001 by Fisher’s Exact test). Within the TCGA data set, we also find increased rates of co-occurrence with Ras GAPs *RASA1* and *RASA3* for noncanonical *KRAS* mutations (Figure 3D). The differences were statistically significant for both of these Ras GAPs (p<0.02 by for *RASA1*, p<0.04 for *RASA3*, both by Fisher’s exact test). Within the larger TCGA data set, mutations were identified in Ras GAPs other than *NF1*, *RASA1*, and *RASA3*. The general trend for increased co-occurrence of noncanonical Ras mutations with mutations in Ras GAPs was again observed (Figure S3). Of note, *RASAL3* also displayed statistically significant rates of increased co-occurrence with noncanonical *KRAS* mutants (p<0.04).

### Noncanonical Ras mutations also co-occur with Ras GEF mutations

Since GAPs and GEFs have opposite effects on Ras, we considered the possibility that partial activation of Ras GEFs will be analogous in some ways to partial loss of Ras GAPs. Partial activation of a Ras GEF could come from a partial increase in upstream signaling or by a GEF mutation. We considered whether such an increase in GEF activation might also influence the effects of noncanonical Ras mutations. To investigate this, we used our mathematical model to simulate the same set of one million computational random Ras mutants within the condition of a partial increase in GEF activity (see Methods and Supplemental Information). Partial GEF activation could also increase the effects of some Ras mutants (Figure 4A). We note that the effects of a GEF mutation or an upstream mutation that results in increased GEF activation would likely have wide range of potential levels of GEF (and resultant Ras) activation. To compare aberrant GEF versus GAP-deficient conditions, we normalized within the model an amount of total basal GEF activity that would result in the same level of basal Ras activation as the NF1-deficient conditions modeled in Figure 1A in the context of wild type Ras.

**Figure 4.**
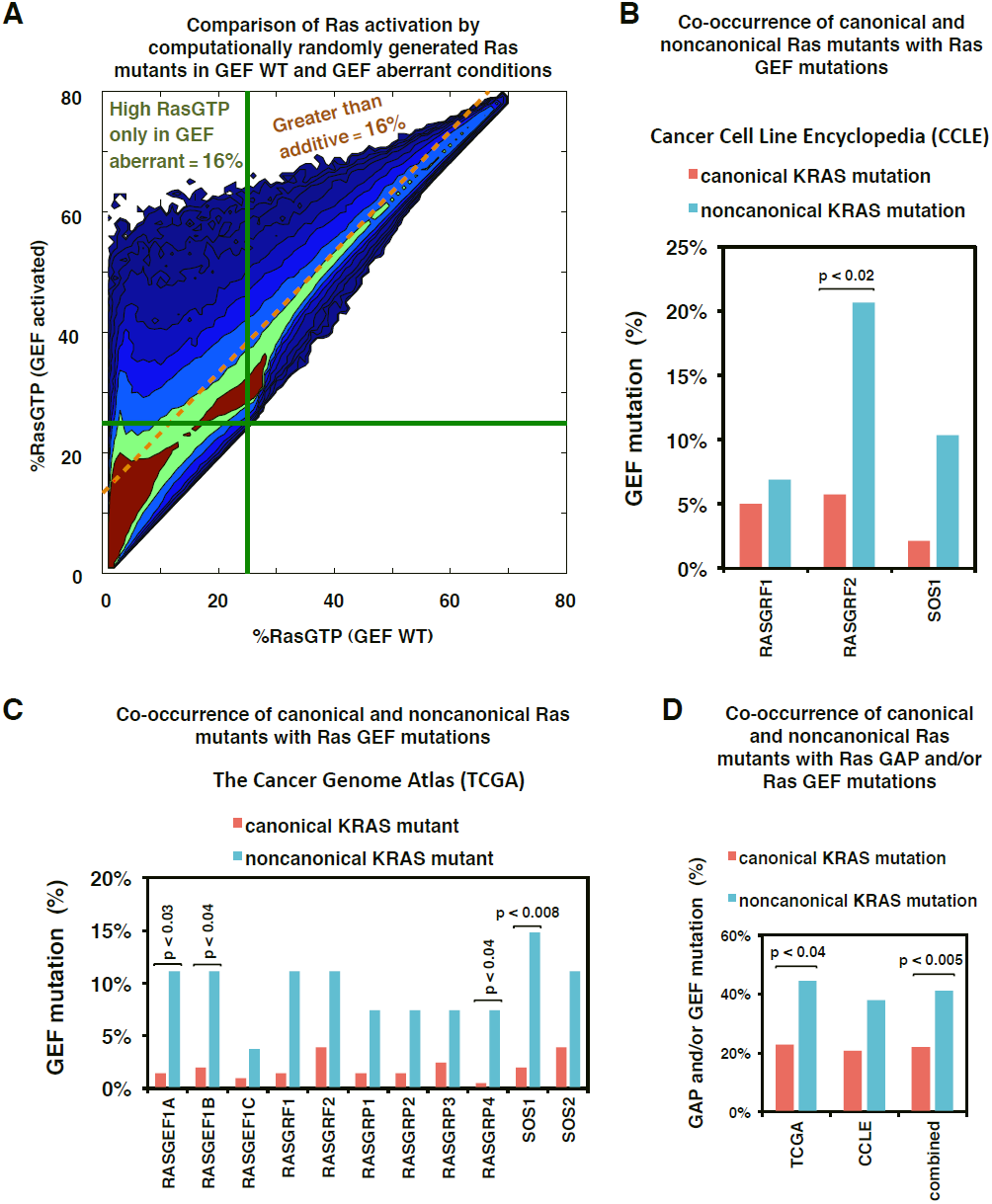
Ras GEF activation can potentiate the effects of a noncanonical Ras mutant. A. One million random mutants were simulated in the WT and GEF activated states, and the resulting levels of RasGTP are plotted for both conditions. Any random Ras mutant falling above the dashed gold line shows a greater net change in percent RasGTP in the GEF activated network compared to the WT network. Green lines indicate 25% total RasGTP.
B. Percentage of canonical and noncanonical *KRAS* mutant cells that co-occur with a Ras GEF mutation within the CCLE dataset (Barretina et al., 2012).
C. Percentage of canonical and noncanonical *KRAS* mutant cancers that co-occur with a Ras GEF mutation within the TCGA datasets (Kandoth et al., 2013).
D. Percentage of canonical and noncanonical *KRAS* mutant samples from the CCLE and TCGA that also harbor at least one GAP or GEF mutant. The p-value for panels B, C, and D were calculated by Fisher’s exact test.

Our computational random mutagenesis found that a modest increase in basal GEF activity could also potentiate the effects of some (but not all) Ras mutants. The proportion of Ras mutants that were strongly activating in GEF aberrant conditions but not in WT conditions (16% of simulated mutants) was similar to that observed for GAP deficiency (15% of simulated mutants). The fraction of mutants that displayed greater than additive behavior with aberrant GEF signaling (16%) was also similar to what was observed for GAP deficiency (13%). Overall, these simulations suggest that mutations that result in increased levels of basal GEF activation may potentiate Ras mutant signals, essentially similar to mutations in NF1 and other Ras GAP proteins.

We next queried the CCLE and TCGA cancer genomic data sets for a hypothesized increased rate of co-occurrence between noncanonical *RAS* mutants and Ras GEF mutants. Within the CCLE data, we found an increased rate of *RASGRF1*, *RASGRF2*, and *SOS1* mutations when noncanonical *KRAS* mutants were present compared to the rate observed when canonical *KRAS* mutants were present (Figure 4B). This difference was statistically significant for *RASGRF2* (p<0.02 by Fisher’s Exact test). Within the TCGA data, we found an increased rate of mutations to all Ras GEFs when noncanonical *KRAS* mutants were present (Figure 4C). This difference was statistically significant for *RASGEF1A* (p<0.03), *RASGEF1B* (p<0.04), *RASGRP4* (p<0.04), and *SOS1* (p<0.008) (all p-values by Fisher’s Exact test).

Since there is an increased rate co-occurrence of either Ras GAP or Ras GEF gene mutation with noncanonical *RAS* mutations, we were curious about the overall rate of co-occurrence, and examined the proportion of canonical and noncanonical *RAS* mutations that co-occur with any Ras GAP or Ras GEF gene mutation. Within the TCGA data set, we found that 23% of canonical *KRAS* mutations co-occurred with a Ras GAP and/or a Ras GEF mutation; remarkably, 44% of noncanonical *KRAS* mutations co-occurred with a Ras GAP and/or a Ras GEF mutation (Figure 4D), p<0.04 by Fisher’s Exact test). We observed a very similar pattern within the CCLE data set; 21% of canonical *KRAS* mutations co-occurred with a Ras GAP and/or a Ras GEF mutation, while 38% of noncanonical *KRAS* mutations co-occurred with a Ras GAP and/or a Ras GEF mutation. When the two datasets were combined, the trend for increased rate of GAP or GEF mutations with noncanonical *KRAS* mutations was highly significant (p<0.005 by Fisher’s Exact test). Overall, the observed increased rate of co-occurrence between noncanonical *RAS* mutations with a Ras GAP or Ras GEF mutation is consistent with our computational model based prediction that Ras GAP and Ras GEF mutations can synergize with some weakly activating Ras mutations to result in a high level of Ras pathway activation.

### Instability of the Ras signaling network in the context of NF1 deficiency

We next set out to investigate possible reasons to explain why GAP deficient and/or GEF activated conditions might augment the effects of some Ras mutations. We hypothesized that the sensitivity of the NF1-WT and NF1-deficient Ras networks might be differentially affected by changes in Ras biochemistry. As a Ras mutation generally results in changes to the rate constants and enzymatic properties of reactions involving Ras, such an analysis could potentially reveal why some Ras mutations may be synergistic with neurofibromin deficiency. Our Ras model includes the five basic processes that regulate the nucleotide binding state of Ras proteins: GAP activity on Ras, GEF activity on Ras, intrinsic GTPase activity, spontaneous nucleotide dissociation and association, and GTP-bound Ras interactions with effector proteins (Figure 5A). Each of these processes is modeled with mass-action kinetics or enzymatic kinetics and described with the parameters listed in Figure 5A.

**Figure 5.**
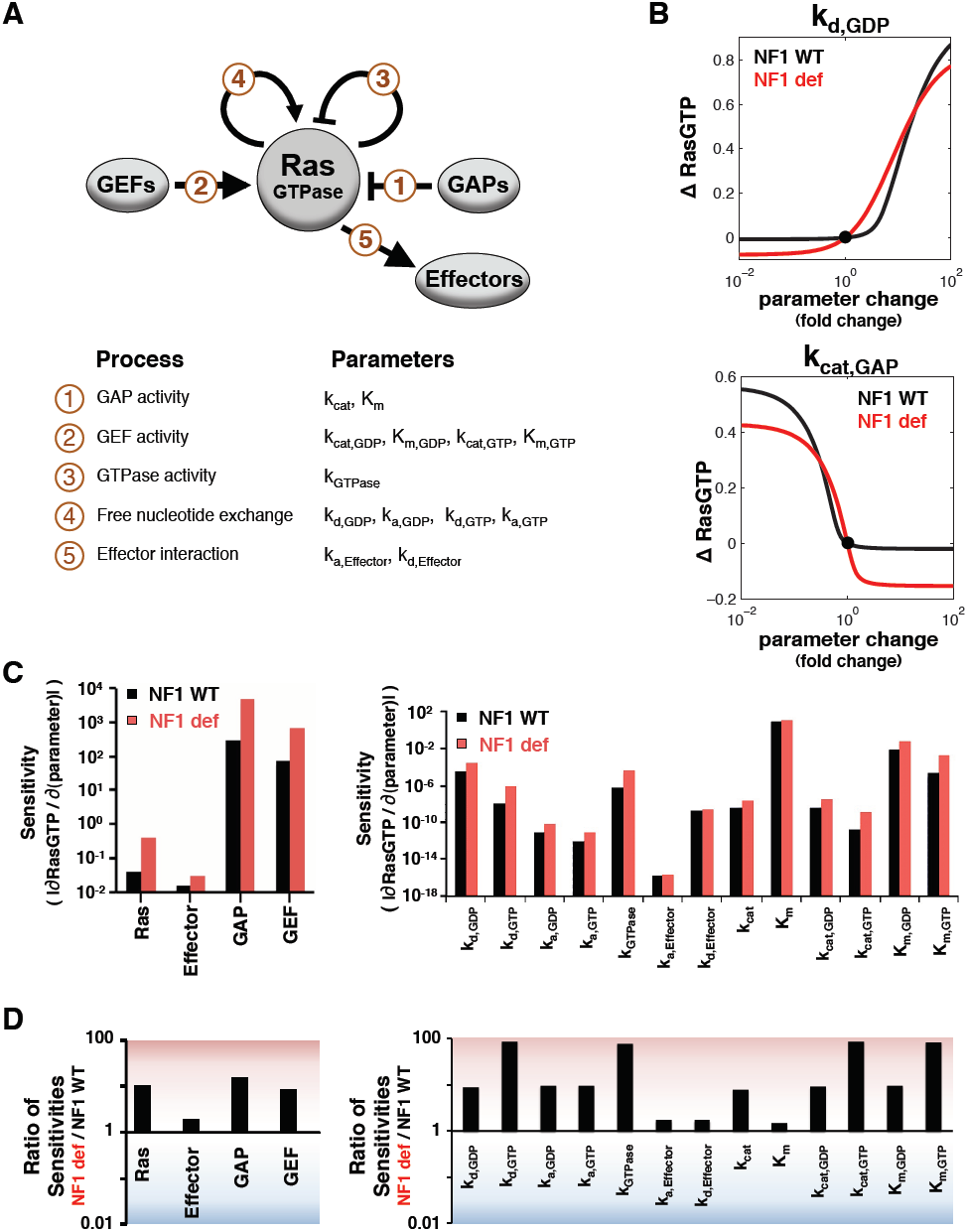
Mathematical model of the Ras network predicts the NF1-deficient Ras network is generally more sensitive to perturbations. A. The key components of the Ras network considered here are Ras, Ras GEFs, Ras GAPs, and Ras effectors. The modeled reactions that influence levels of RasGTP and the biochemical parameters for these reactions are indicated.
B. Net changes in RasGTP levels from the corresponding, baseline, steady-state for a range of fold-changes in Ras network rate constants and enzymatic parameters for both NF1-WT and NF1-deficient conditions. Values for the dissociation of GDP from Ras (k_d,GDP_) and the kcat of the Ras GAP reaction on Ras GTP (k_cat,GAP_) are presented, with all parameters presented in Figures S4 and S5. **Δ**RasGTP levels are normalized to the total amount of Ras. The data suggests that the NF1-deficient state will experience a larger change in RasGTP for a small change in protein expression.
C. The magnitude of the immediate rate of change in RasGTP for a change in rate constant or enzymatic parameter is a measure of the sensitivity of the Ras network to that parameter.
D. The ratio of the sensitivities determined in C. This suggests that the NF1-deficient state will experience a larger magnitude change in RasGTP for a small change in any parameter.

We considered the magnitude of change in total RasGTP that would result for a wide range of changes in each parameter when it occurs in NF1-WT and in NF1-deficient conditions (Figures 5B **and** S4-S5). Qualitatively, the relationships were similar for NF1-WT and deficient networks when we considered the amount of RasGTP that results from a change in a single reaction parameter. It was notable, however, that the magnitude of change in RasGTP for a small change in a parameter was always greater for the NF1-deficient network (Figures 5B **and** S4-S5). This was true for all 17 of the model’s biochemical properties. Such a distribution would be highly unlikely by chance (P<1.6 × 10^−5^ by the two-tail exact binomial test). We note that we focus here on the absolute increase in RasGTP from the respective NF1-WT and NF1-deficient conditions rather than the relative change in RasGTP because it appears that the total amount of Ras pathway activation, and not the relative change from each mutation, represents an important feature in cancer cell signaling (Donovan et al., 2002; Shapira et al., 2007).

To further investigate the relevance of this observation, we analytically solved for the change in RasGTP for a change in parameter 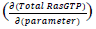 for both NF1-WT and NF1-deficient networks (i.e. the slope of the curves relating RasGTP to parameter values taken through the point corresponding to the baseline Ras network parameters). Analytic calculations were based on the same set of reaction equations that were used in the simulation model. These reaction equations were algebraically reduced to an expression that relates the levels of RasGTP to each parameter, and this equation was then implicitly differentiated. The resulting expression was then numerically evaluated at the steady-state level of RasGTP with the model parameters. The values of the slopes determined from this analytic approach matched the values derived from numerical simulations. As inferred from the graphical relationship between RasGTP and changes in model parameters, the analytical approach revealed that the net change in RasGTP induced by a small change in a reaction rate constant or enzymatic parameter is higher in the NF1-deficient network (Figure 5C,D). Indeed, the NF1-deficient network was often 10-100 times more sensitive to the same change in a network reaction rate constant and/or protein concentration. Thus, any small perturbation to the Ras network should cause a larger magnitude change in Ras signal if the network is also NF1-deficient.

We also noted that the increased magnitude change in RasGTP does not occur for some large changes to reaction rate constants (seen in Figures 5B and S4-S5 for several parameters). This is likely because the NF1-deficient network has a higher proportion of total Ras in the RasGTP form, and a smaller net change is needed to reach saturation as RasGTP. However, the level of constitutive Ras pathway signaling associated with disease causing mutations appears to be far less than saturating (Donovan et al., 2002). Thus, the increased magnitude change within NF1-deficient conditions that occurs from an already elevated baseline is quite relevant to cancer progression.

We further analyzed our model to evaluate the robustness of the prediction that neurofibromin deficiency amplifies the effects of perturbations within the Ras network (see Supplemental Information). We found that our model predictions were robust to the level of GAP deficiency considered (Figure S6), and to the concentrations of Ras network proteins modeled (Figure S7). We also performed a similar analysis for increased basal Ras GEF activity (such as what might follow from an upstream mutation, or from a Ras GEF mutation) (see Supplemental Information). We found that increased basal GEF activity also makes the Ras network more sensitive to perturbations (Figure S8). That is, increased sensitivity to perturbations should be a more general feature of cells harboring Ras GAP and Ras GEF mutations.

### General increase in co-occurrence of Ras pathway mutations in cancer genomes

We hypothesized that the increased sensitivity to perturbation should also result in an increased co-occurrence of *NF1* mutations with mutations to the genes of Ras network proteins (GEFs, GAPs, and effectors). Within the CCLE data the frequency of Ras network mutations appeared higher for *NF1* mutant cells than for *NF1* wild-type cells (Figure 6A). The increased co-occurrence was statistically significant for Ras effectors *PIK3CA* and *ARAF*, as well as for upstream *EGFR* mutations, and for several Ras GAPs and Ras GEFs. Of note, the increased co-occurrence of *all* (i.e. canonical and noncanonical) *KRAS*, *NRAS*, and *HRAS* mutations was not statistically significant, consistent with neurofibromin deficiency having only a small effect on RasGTP for the more common canonical Ras mutants (Figure 1A). We also note that rates of *TP53* mutations were not appreciably different between *NF1* mutant and *NF1* WT samples (Figure 6B).

**Figure 6.**
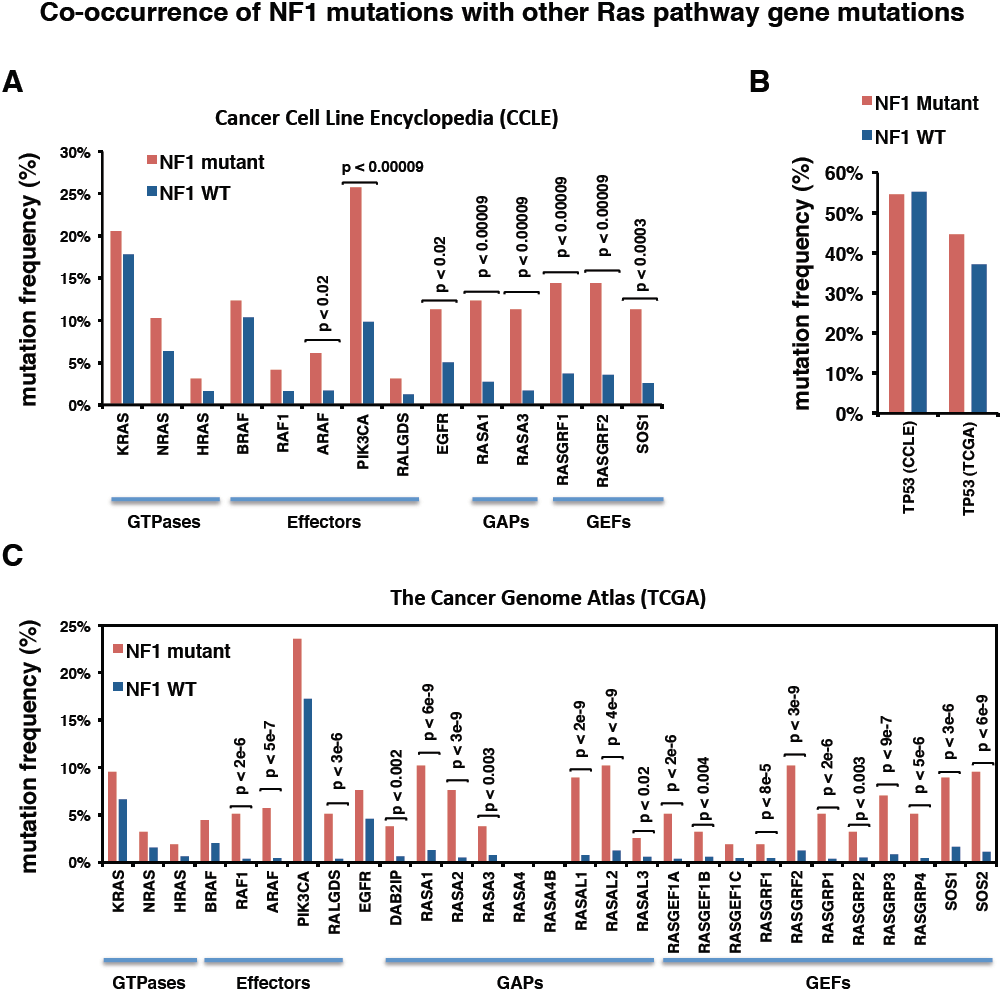
Analysis of cancer genomic data finds the predicted increase in co-occurrence between *NF1* and less commonly mutated Ras genes. A. Mutation frequency for Ras network genes within *NF1* mutant and *NF1* WT subsets of the CCLE dataset (Barretina et al., 2012).
B. Mutation frequency for *TP53* in both *NF1* mutant and *NF1*WT samples in the CCLE and TCGA datasets.
C. Mutation frequency for Ras network genes within *NF1* mutant and *NF1* WT subsets of the TCGA dataset (Kandoth et al., 2013). The p-value for all panels is by Fisher’s exact test.

We similarly analyzed the larger TCGA data set. We again found a trend for increased co-occurrence of less commonly mutated Ras pathway genes along with *NF1* mutations (Figure 6C). Of note, we found increased rates of co-occurrence between *NF1* mutations and noncanonical Ras effector mutations, *RAF1* and *ARAF* and *RALGDS*. With the much larger sample size, the differences were statistically significant for these Ras effectors (p<2×10^−6^, p<5×10^−7^, p<3×10^−6^, respectively, by Fisher’s exact test). This increased level of co-occurrence suggests that these less commonly mutated Ras pathway genes do contribute to the cancer phenotype when they are found with an NF1 mutation.

Additionally, co-mutations with other Ras GAPs and Ras GEFs were quite enriched, which is consistent with the predicted synergy between Ras GAP and Ras GEF mutations (e.g., Figure 5C,D). Co-mutations between *NF1* and other Ras GAPs were statistically significant for *NF1* with *DAB2IP* (p<0.002), *RASA1* (p<6×10^−9^), *RASA2* (p<3×10^−9^), *RASA3* (p<0.003), *RASAL1* (p<2×10^−9^), *RASAL2* (p<3×10^−9^), and *RASAL3* (p<0.02). Co-mutations between *NF1* and Ras GEFs were statistically significant for *RASGEF1A* (p<2×10^−6^), *RASGEF1B* (p<0.004), *RASGRF1* (p<8×10^−5^), *RASGRF2* (p<3×10^−9^), *RASGRP1* (p<2×10^−6^), *RASGRP2* (p<0.003), *RASGRP3* (p<9×10^−7^), *RASGRP4* (p<5×10^−6^), *SOS1* (p<3×10^−6^), and SOS2 (p<6×10^−9^). Within the TCGA dataset, as was the case in the CCLE data set, there was no significant difference in the rate of *TP53* mutations in *NF1* mutant and *NF1* WT samples (Figure 6B).

Overall, these findings support the model-based hypothesis that neurofibromin deficiency results in a system that is more sensitive to mutations elsewhere in the Ras pathway. That the increased co-occurrences are observable in cancer sample genomic data may further suggest that these co-occurrences have effects sufficient to confer a competitive advantage to the cell that first acquires the combination of mutations.

## DISCUSSION

It has been argued that better methods are needed for analyzing the relationship between mutations and the perturbed cell signaling networks that drive cancer and may also provide therapeutic targets (Yaffe, 2013). We have here presented here the use of mass-action modeling to uncover previously unappreciated relationships between pairs of mutations and found evidence for increased co-occurrence of these pairs of mutations in existing cancer genomic data sets. Our model was based on the traditional understanding of Ras network biochemistry and the available, quantitative measurements that characterize Ras network biochemistry. Model predictions have been prospectively validated experimentally in mammalian cells. Importantly, insights gained from these computational and experimental studies generated specific hypotheses that we could test in existing, large, cancer genomic data sets. Such an approach should be more generally applicable to other signaling networks.

It has become increasingly clear that the strongly activating Ras mutants in codons 12, 13, and 61 are not the only significant Ras mutants found in human cancers and disease. Noncanonical, weakly activating Ras mutants have been linked to Noonan syndrome (Schubbert et al., 2006). Comprehensive screening of the *RAS* genes in myeloid leukemia samples found noncanonical *RAS* mutants in 2% of all leukemia samples, representing 14% of all *RAS* mutants observed (Tyner et al., 2009). Additionally, screening for *KRAS* mutations in colorectal cancer found noncanonical Ras mutants in 9% of cancer samples, representing 26% of all *KRAS* mutants observed (Smith et al., 2010). Therefore, our computational and experimental observation that neurofibromin deficiency can potentiate the effects of noncanonical Ras mutants is very relevant to cancer development.

The finding of an increased co-occurrence of mutations within the same pathway is surprising in that conventional wisdom holds that mutations within the same pathway do not co-occur (Thomas et al., 2007; Yeang et al., 2008). However, conventional wisdom is based on the behavior of the strongly activating, canonical mutations, which do not co-occur. Indeed, our modeling suggests that combinations of mutations involving a strongly activating canonical mutation have a less-than-additive total response and would therefore not be expected to acquire a significant competitive advantage (Figure 1A). Therefore, our modeling is consistent with the lack of co-occurrence between strongly activating canonical mutations. An unbiased approach at combination discovery would require a very large sample size because of the very large number of potential combinations. The evaluation of mutation co-occurrence without a prior hypothesis is further confounded by the smaller number of mutations observed for atypically mutated gene compared to the number of mutations observed for canonical cancer driver genes. With a very specific hypothesis derived here from mathematical modeling, a smaller number of sequenced cancer genomes enables discovery of enriched combinations of mutations and the finding that conventional wisdom does not apply to noncanonical mutations.

The finding that the Ras network with a weakly activating mutation is generally more sensitive to subsequent perturbations has important implications in cancer development. The cancer phenotype results from the acquisition of multiple somatic mutations that ultimately result in altered levels of protein expression or the expression of a protein with altered biochemistry (Stratton et al., 2009). The computational model-based predictions and experimental work presented here suggest that the number of biochemical perturbations (and causal genetic aberrations) with a large effect on Ras pathway signal increases in the context of neurofibromin deficiency. This would expand the number of potential “driver genes” that promote cancer in the neurofibromin-deficient context. The net rate of acquiring a cancer-promoting mutation is proportional to both the rate of mutation and the proportion of mutations that offer a selective advantage to the cell. Genetic instability is a common feature of cancer that results in an increased rate of mutations that could promote tumorigenesis (Beckman and Loeb, 2005). Our work suggests an alternative, yet complementary mechanism of “network instability” that results in an increased net rate of acquiring cancer-promoting mutations through the acquisition of a state where a greater proportion of mutations would offer a selective advantage.

## EXPERIMENTAL PROCEDURES

### Mathematical model

The mathematical model of the Ras signaling network that is the basis of this work has been previously described extensively (Stites and Ravichandran, 2012a; Stites et al., 2007). The model was developed, simulated, and analyzed in MATLAB v7.11.0.584 (R2010b) (MathWorks). Algebraic manipulations and numerical evaluations of the algebraic equations were performed in Mathematica 8 (Wolfram). NF1 deficiency was modeled by decreasing the concentration of total GAP in the model. For NF1 WT conditions, we used the full concentration of Ras GAP identified in our original Ras network model, and we used 50% of this value to model NF1-deficient conditions. An increase in basally active GEF concentration that would result in the same level of RasGTP as a 50% decrease in basally active GAP was used to model aberrant GEF activation. Simulations and/or analytical calculations were used to determine model predicted levels of RasGTP.

The change in RasGTP for a change in a model parameter was determined with simulations and/or with analytical calculations. For simulations, a single parameter was adjusted by ± 0.1% of its value in the WT network, and the resultant level of RasGTP was determined with model simulations. The difference in RasGTP, or ΔRasGTP, was determined from the two resulting levels of steady-state RasGTP. This was done for WT and NF1/GAP-deficient networks. The ratio of ΔRasGTP for the NF1-deficient network to ΔRasGTP for the WT network was used as a measure of relative sensitivity to a change in a single parameter value. Analytical calculations were performed using the same set of reactions and conservation laws for total Ras and total effector. Algebraic steady-state solutions were obtained, and further algebraic manipulation reduced these equations to a single expression that implicitly relates steady-state levels of RasGTP to each parameter, *G*(RasGTP,k) = 0 where k indicates all of the parameters of the model (including concentrations, rate constants, and enzymatic parameters. To determine 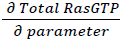 for each parameter, we use the implicit function theorem, with 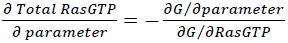

Each expression was then numerically evaluated with the model parameter values. This was done for WT (100% basal GAP present) and NF1-deficient (50% basal GAP present) conditions. The ratio of 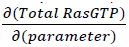 for the NF1-deficient network to the 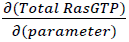 for the WT network was used as a measure of relative sensitivity to a change in a single parameter value, just as when these changes were determined with simulations.

Computational random Ras mutants were generated by obtaining a random set of factors (a_i_ with i=1:12) with which to vary each of the twelve independent RasWT reaction parameters. Each random mutant was generated by first creating a set of 12 random numbers from a random number distribution with a mean of zero and standard deviation of 1 (x_i_ with i = 1:12). This random number, x_i_, was transformed into the corresponding parameter multiplication factor, a_i_=10^x^. These were then applied to twelve free reaction parameters (k_mut,i_ = a_i_ k_WT,i_). The remaining parameter, (k_catGTP_) was calculated from the other parameters to ensure thermodynamically consistent parameters for nucleotide exchange (whether GEF-mediated or free nucleotide exchange).

### Cell transfection

Immortalized *Nf1*+/+, *Nf1*+/-, and *Nf1*-/- MEF cells were obtained from Dr. Reuven Stein at the George S. Wise Faculty of Life Sciences, Tel Aviv, Israel. Cells were maintained as described previously (Shapira et al., 2007). The plasmids encoding 3xHA-tagged RasWT and RasG12V were obtained from Missouri S&T cDNA Resource Center. Ras mutants RasG12D and RasF28L were generated using the QuikChange mutagenesis kit (Stratagene) and confirmed by sequencing. V5 tagged NF1-GRD was provided by Dr. David Largaespada at the University of Minnesota, Minneapolis, MN. MEF cells were plated on six-well plates and transiently transfected with 3 µg of plasmid constructs using Lipofectamine 2000 and according to manufacturer’s instructions (Invitrogen). After 4 hours, fresh growth medium was added for 6 hours. Cells were starved for 12 hours in DMEM 0.5% fetal calf serum (FCS) and then harvested for cytometry analysis.

### Immunoblotting

MEF cells were subjected to lysis in a buffer containing 50 mM Tris (pH 7.6), 150 mM NaCl, 1 mM EDTA, 1% Triton X-100 10 mM sodium pyrophosphate, 10 mM sodium fluoride, 1 mM sodium orthovanadate, and protease inhibitors (Calbiochem). Total cell lysates were separated by 5-20% gradient SDS-PAGE and transferred to PVDF membrane. Membranes were blocked with 5% milk in 1% tween-20 Tris-buffered saline buffer. Proteins were detected by antibodies against NF1 (Upstate Biotechnology, Millipore), V5-tag (Millipore), p120 RasGAP (BD biosciences), Erk1/2 (Cell signaling Tech), phospho-Erk1/2 (Cell Signaling Tech), HA-tag (Cell Signaling Tech). All immunoblots were developed using enhanced chemiluminescence (Pierce, Rockford, IL).

### Quantitative PCR

Total RNAs were extracted from MEF cells with a QIAshredder and RNeasy kit (Qiagen). The SuperScript III kit (Invitrogen) was used for reverse transcription. TaqMan Gene Expression assays for *Rasa1*, *Rasa4*, *DAB2IP*, and *Nf1* (Applied Biosystems) were used for quantitative PCR. Samples were amplified in duplicate and target transcripts were normalized to *Hprt1* mRNA as a housekeeping gene. The relative expression of each target gene was calculated by the comparative cycling method with StepOne v2.1 software (Applied Biosystems). The standard deviation was calculated after normalization of multiple experiments.

### Flow cytometry

MEF cell staining for phospho-Erk1/2 (referred to as phoshpo-ERK or pERK) and HA-tagged Ras or other proteins of interest was performed as described previously (Stites et al., 2007). Data were acquired on a FACS CantoII (Becton Dickinson). Cells were gated based on the intensity of the HA signal to define “high” and “low” HA expressing populations. To assess cell proliferation and allow cell cycle synchronization, MEF cells were starved for 12 hours in DMEM 0.5% FCS and stained with CellTrace™ Violet proliferation kit, according to manufacturer’s instructions (Invitrogen). Stained cells were then harvested in a time course and their state of proliferation analyzed by cytometry.

### Cancer genome analysis

Mutations for the genes of Ras network proteins (*ARAF*, *BRAF*, *DAB2IP*, *EGFR*, *HRAS*, *KRAS*, *NF1*, *NRAS*, *PIK3CA*, *RAF1*, *RALGDS*, *RASA1*, *RASA2*, *RASA3*, *RASA4*, *RASA4B*, *RASAL1*, *RASAL2*, *RASAL3*, *RASGEF1A*, *RASGEF1B*, *RASGEF1C*, *RASGRF1*, *RASGRF2*, *RASGRP1*, *RASGRP2*, *RASGRP3*, *RASGRP4*, *SOS1*, and *SOS2*) and for *TP53* were obtained from the CCLE portal (http://www.broadinstitute.org/ccle) and/or from published cancer genome publications (Barretina et al., 2012; Kandoth et al., 2013). Exonic missense and nonsense mutation, and exonic insertions and deletions were considered. Coincident mutations were counted as the number of distinct samples with at least one mutation in specified genes. Fisher’s exact test was used to determine p-values for co-occurring mutations, and calculations were performed in R version 2.13.0.

## SUPPLEMENTAL INFORMATION

Supplemental information includes eight figures.

## ACKNOWLEDGMENTS

We are grateful to David Largaespada and Reuven Stein for sharing reagents, as well as to the members of the Ravichandran laboratory and to the University of Virginia Computational Systems Biology community for their helpful comments and conversations. This work was supported by DOD Award W81XWH-09-1-0087 to K.S.R. and by NIGMS award GM55761 to K.S.R. E.C.S. was partially funded by the TGen Foundation as the Randy Pausch Scholar. K.S.R. is Bill Benter Senior Fellow of the American Asthma Foundation and thanks the support via the Harrison Professorship.

---

**Figure S1.**
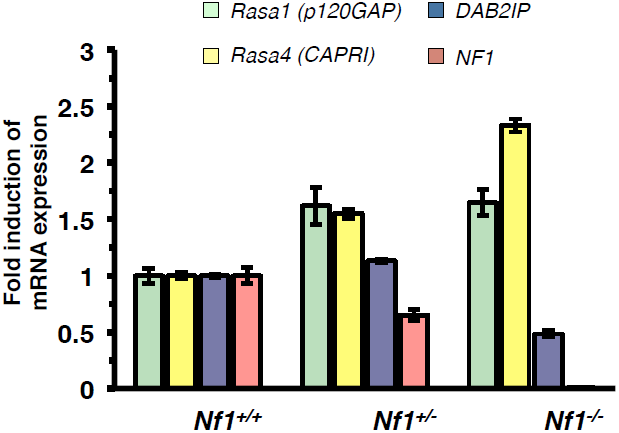
The expression of Ras GAPs in Nf1-deficient mouse embryo fibroblast cells (MEFs) MEFs of the *Nf1*+/+, *Nf1*+/-, and *Nf1*-/- genotype were analyzed by qPCR for Ras GAP genes *Rasa1* (p120GAP), *Rasa4* (CAPRI), *DAB2IP*, and *Nf1*. Error bars represent variation from three independent experiments from three different RNA extractions/preparations.

**Figure S2.**
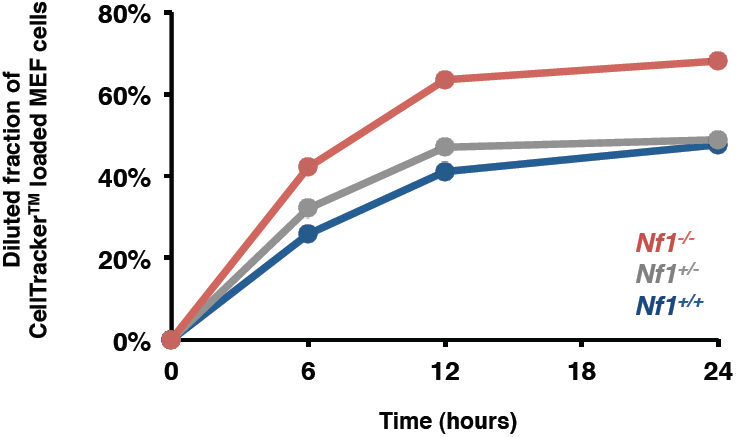
Comparison of proliferation by MEFs with varying levels of neurofibromin expression. Proliferation assay for Nf1+/+, Nf1+/-, and Nf1-/- MEFs.

**Figure S3.**
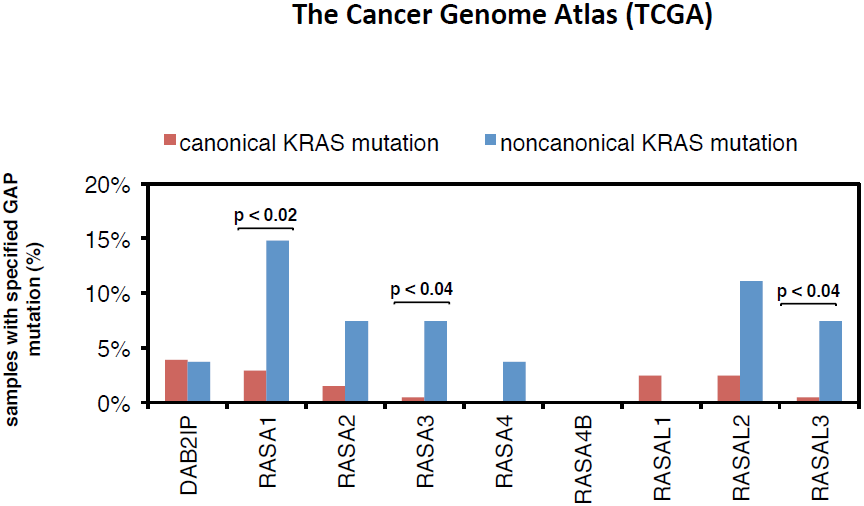
Co-occurrence of noncanonical Ras mutants with mutations of Ras GAPs within the TCGA data. Percentages of canonical and noncanonical K-Ras mutants that co-occur with a Ras GAP mutation within the TCGA dataset are shown. The p-value for all panels is by Fisher’s exact test. Data for *RASA1* and *RASA3* are also included in Figure 3D.

**Figure S4.**
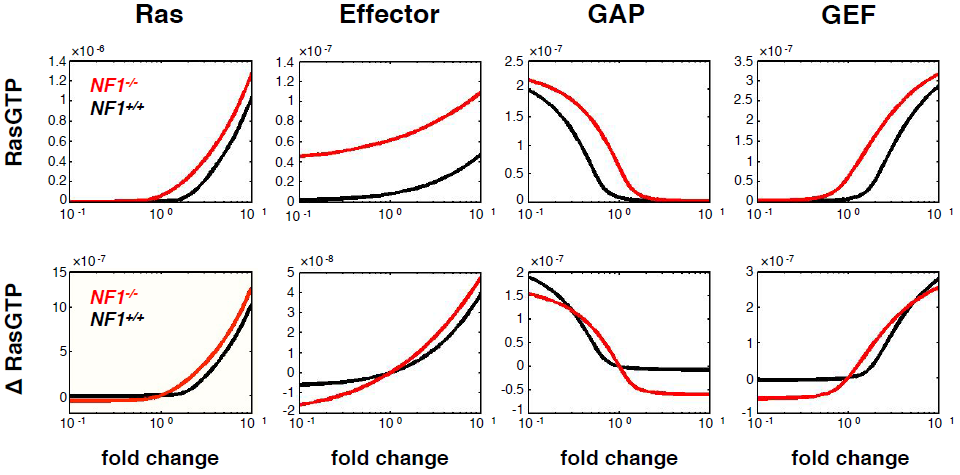
Net change in RasGTP for a change in the indicated Ras network concentration parameter. RasGTP levels (M) and net changes in RasGTP levels are determined through model simulations when concentration parameters are varied in WT and NF1-deficient networks. Parameters are varied in terms of fold change from the baseline value of the parameter. **Δ**RasGTP levels are normalized to the total amount of Ras.

**Figure S5.**
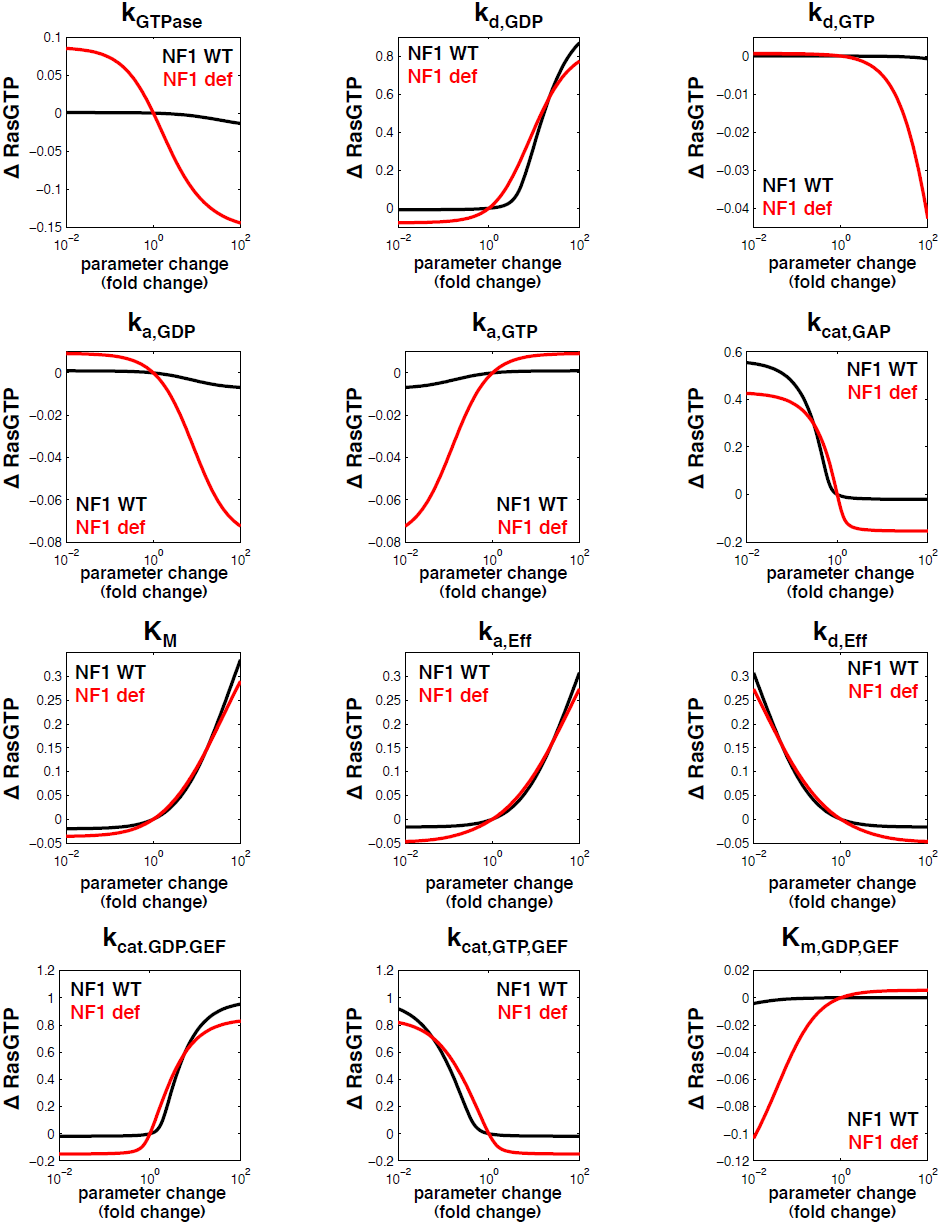
Net changes in RasGTP for a change in the indicated Ras network reaction parameter. Net changes in RasGTP levels determined through model simulations when the reaction parameters are varied in NF1-WT and NF1-deficient networks. Parameters are varied in terms of fold change from the baseline value of the parameter. **Δ**RasGTP levels are normalized to the total amount of Ras.

**Figure S6.**
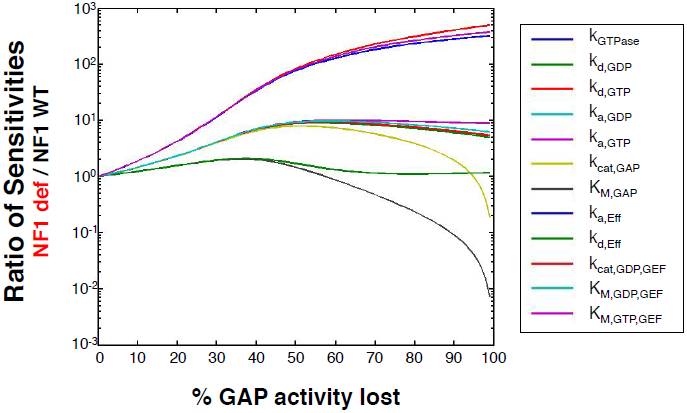
The proportion of total basal GAP activity lost influences the magnitude and sensitivity of the GAP-deficient network. The ratio of sensitivities (i.e. the change in RasGTP for a small change in each parameter) between GAP-deficient (e.g. NF1-deficient) and GAP-WT networks was determined for varying levels of basal GAP activity lost.

**Figure S7.**
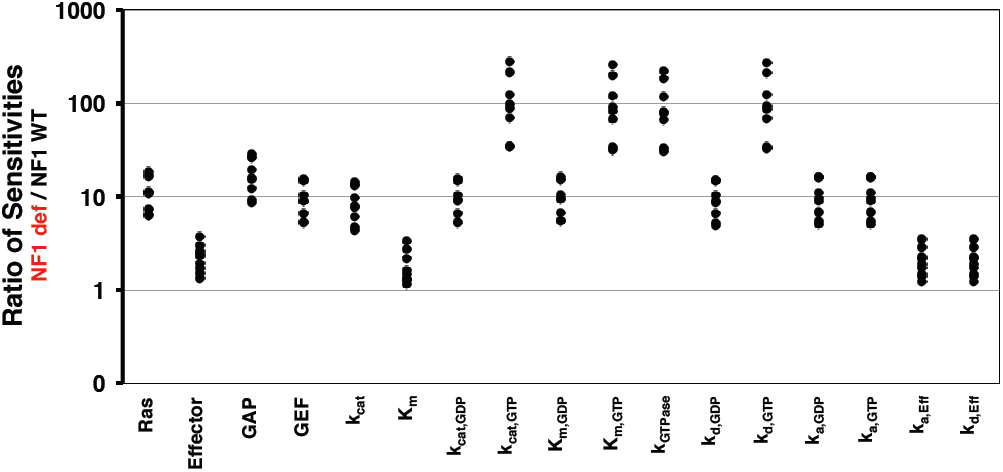
The model-based predictions that NF1-deficient networks are more sensitive to perturbation are robust to changes in the basal protein concentrations. Nine concentration sets, which were previously used to assess the robustness of the Ras network model to the changes in specific concentrations of network proteins, were used here to assess predictions about neurofibromin deficiency. The same nine concentration sets were applied to the analysis presented in Figure 5D. Increased sensitivity of the NF1-deficient network across all model parameters is a robust result that is not dependent upon specific network protein concentrations used. Each data point represents the ratio of sensitivities between NF1-deficient and NF1-WT networks for the specified parameter for one of the nine concentration sets.

**Figure S8.**
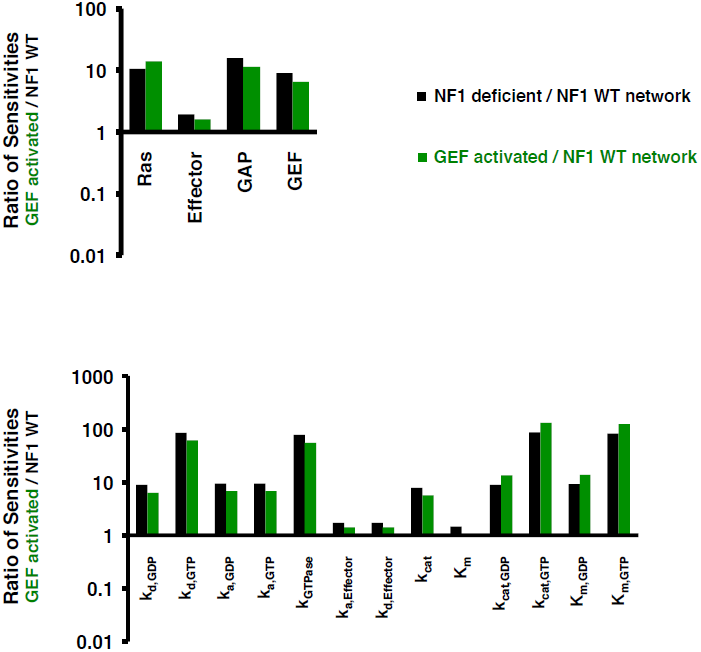
Ras networks with increased basal GEF activity are more sensitive to perturbation. Calculations of the sensitivity of RasGTP levels to changes in model parameters were performed for networks with increased basal GEF activity and compared to the wild-type network, similar to what was done in Figure 5D for neurofibromin deficiency. The ratio of sensitivities between the GEF activated and the NF1-WT network is presented (green). The ratio of sensitivities between the NF1-deficient network and the NF1-WT network is reproduced here for comparison (black).

